# *In Silico* Design of siRNAs Targeting Existing and Future Respiratory Viruses with VirusSi

**DOI:** 10.1101/2020.08.13.250076

**Authors:** Dingyao Zhang, Jun Lu

**Author notes:** Correspondence should be addressed to: Jun Lu, 10 Amistad Street, Room 237C, New Haven, CT 06520.

## Abstract

The COVID-19 pandemic has exposed global inadequacies in therapeutic options against both the COVID-19-causing SARS-CoV-2 virus and other newly emerged respiratory viruses. In this study, we present the VirusSi computational pipeline, which facilitates the rational design of siRNAs to target existing and future respiratory viruses. Mode A of VirusSi designs siRNAs against an existing virus, incorporating considerations on siRNA properties, off-target effects, viral RNA structure and viral mutations. It designs multiple siRNAs out of which the top candidate targets >99% of SARS-CoV-2 strains, and the combination of the top four siRNAs is predicted to target all SARS-CoV-2 strains. Additionally, we develop Greedy Algorithm with Redundancy (GAR) and Similarity-weighted Greedy Algorithm with Redundancy (SGAR) to support the Mode B of VirusSi, which pre-designs siRNAs against future emerging viruses based on existing viral sequences. Time-simulations using known coronavirus genomes as early as 10 years prior to the COVID-19 outbreak show that at least three SARS-CoV-2-targeting siRNAs are among the top 30 pre-designed siRNAs. Before-the-outbreak pre-design is also possible against the MERS-CoV virus and the 2009-H1N1 swine flu virus. Our data support the feasibility of pre-designing anti-viral siRNA therapeutics prior to viral outbreaks. We propose the development of a collection of pre-designed, safety-tested, and off-the-shelf siRNAs that could accelerate responses toward future viral diseases.

## Introduction

The COVID-19 pandemic has highlighted the challenges posed by emerging respiratory viruses to human societies. Other than quarantine and social distancing, existing off-the-shelf anti-viral drugs often do not work well against these new viruses. How to combat COVID-19 and future emerging respiratory viruses thus need to be urgently addressed.

SARS-CoV-2, the COVID-19-causing coronavirus, belongs to a family of positive-sense and single-strand RNA viruses(1–3). This virus family is notorious for its high mutation rates, which suggests that immunity triggered by either a vaccine or a previous infection may not be long-lasting. Targeting the SARS-CoV-2 genome using small interfering RNAs (siRNAs) or CRISPR can be an alternative therapeutic approach. This is because a mixture of multiple siRNAs or CRISPR RNAs, targeting different regions of the viral genome, could theoretically be delivered into susceptible tissues such as lung epithelium. Cleavage of viral RNA by siRNA or CRISPR could thus reduce the viral load and limit further infection(4–6). Recently, Timothy et al. reported a proof-of-concept Cas13d system for degrading viral RNAs (7), in which the intriguing idea of designing crRNAs that target multiple respiratory viruses has been explored (7). However, the design of crRNAs in that study uses SARS-CoV-2 sequence as an input, which does not test the feasibility toward preparing for a future viral outbreak. Compared to CRISPR, there are several advantages of using siRNAs as anti-viral therapies. The siRNA approach avoids the challenge of delivering large Cas gene products into cells *in vivo*. Efficacy of a cocktail of multiple siRNAs against the EBOLA virus has been demonstrated successful *in vivo* in primate models(8). Additionally, pulmonary delivery of siRNAs is possible to knock down target genes in animal lung(9–12), and has been used to limit respiratory viruses in animal models(13). A recent preprint further shows the feasibility of intranasal administration of siRNAs to reduce SARS-CoV-2 viral load and symptoms in Syrian hamster and *Rhesus macaque* models (14).

The exciting idea of designing siRNAs against SARS-CoV-2 has been explored in several recent reports(14–19). These studies have used SARS-CoV-2 sequences to design siRNAs against SARS-CoV-2 itself, rather than exploring the possibility of designing siRNAs to guard against future viruses. Furthermore, there are several technical limitations in these existing studies. First, limited numbers (maximum of 143) of SARS-CoV-2 genomes have been used, which could make some previously designed siRNAs fall into genetically variable regions of the viral genome. Second, off-target effects of siRNAs have not been always considered, with potential unintended regulation of endogenous genes by designed siRNAs. Third, given the ever-evolving mutational landscape of SARS-CoV-2 genomes and potential emergence of future respiratory viruses, a computational tool that can design siRNAs against both existing and future viruses is urgently needed.

In this study, we present VirusSi, a computational pipeline that can either rationally design siRNAs against many strains of a single virus, or pre-design siRNAs against future emerging viruses based on existing viral sequences. We demonstrate the feasibility of the pre-design approach and propose a model in which pre-designed siRNAs can undergo testing and manufacturing multiple years prior to a viral outbreak to help accelerate response toward future respiratory viruses. We also used the VirusSi to design multiple siRNAs against SARS-CoV-2, which can be tested in wet-lab settings for efficacy and safety.

## Results

As a first step toward designing siRNAs against SARS-CoV-2 and creating the VirusSi pipeline (see **Figure 1** for flow charts of VirusSi), we established two sets of rules for candidate siRNAs (**Supplementary Table S1**). The first rule set was modified from those in the GPP Web Portal for short hairpin RNA designs. In addition, the VirusSi pipeline provides a second rule set based on those used in the RNAxs program (20) (see Methods) that users can choose from. For the first rule set, we kept several criteria used in GPP Web Portal: excluding any siRNA candidate ending with TT, excluding any candidate containing a run of four of the same base in a row, selecting siRNA candidates with proper AT percentage (25% to 60%), excluding any candidate with a run of 7 C/G bases, excluding any candidate containing ambiguous bases, and giving precedence to candidates with weaker base-pairing at positions 15-20. Additionally, we added the requirement for the first base of the functional siRNA strand to be a U. This is because the crystal structure of human Argonaute 2 (AGO2), the core component of the siRNA machinery, has revealed a preference of binding to small RNAs with a first base of U or A(21, 22), and because the majority of endogenous AGO2-bound RNA, namely miRNAs, start with a U(23). Furthermore, we eliminated several rules in the GPP Web Portal: the exclusion of restriction site which only applies to the cloning of shRNAs, the consideration of potential stem-loop structure in target site because we incorporated another strategy for the influence of target RNA structure (see below).

**Figure 1.**
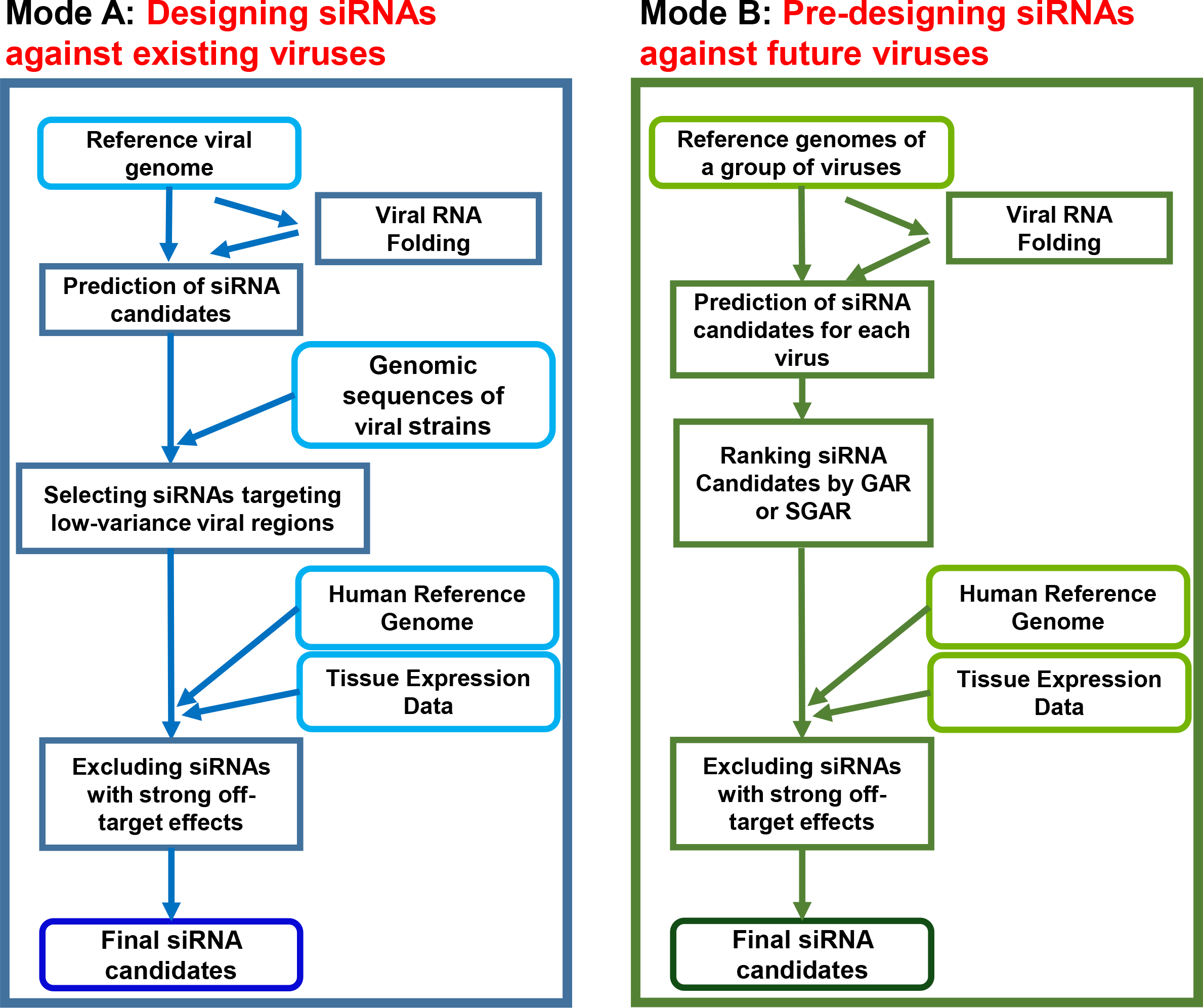
The VirusSi Workflow. **(A)** Mode A of VirusSi, which designs siRNAs against an existing virus of interest, utilizing the reference genome as well as genomic sequences of multiple strains of the virus as inputs. Additionally, human reference genome and the transcriptome for tissue(s) of interest were also used to weed off siRNAs with strong off-target effects. **(B)** Mode B of VirusSi, which designs siRNAs against future emerging viruses, utilizing the existing genomes of a class of viruses (e.g. human and bat coronaviruses) as inputs. This mode pre-designs siRNAs, with ranking of siRNAs by the GAR and SGAR algorithms, so that a small number of top candidates could be tested for safety and efficacy in preparation against future viral outbreaks.

Based on the above siRNA design rules, we predicted candidate siRNA targeting sites according to the reference genome of SARS-CoV-2. We excluded siRNAs that target high-probability double-stranded RNA regions in the SARS-CoV-2 genome, by evaluating the *in silico* folding structure of SARS-CoV-2 and the base-pairing probability for each siRNA scanning window in the viral genome. Target regions that are highly structured were then filtered out (see Methods for details), resulting in 4482 siRNA candidates. These candidates were distributed across all the genes of SARS-CoV-2 (**Figure 2A-B**).

**Figure 2.**
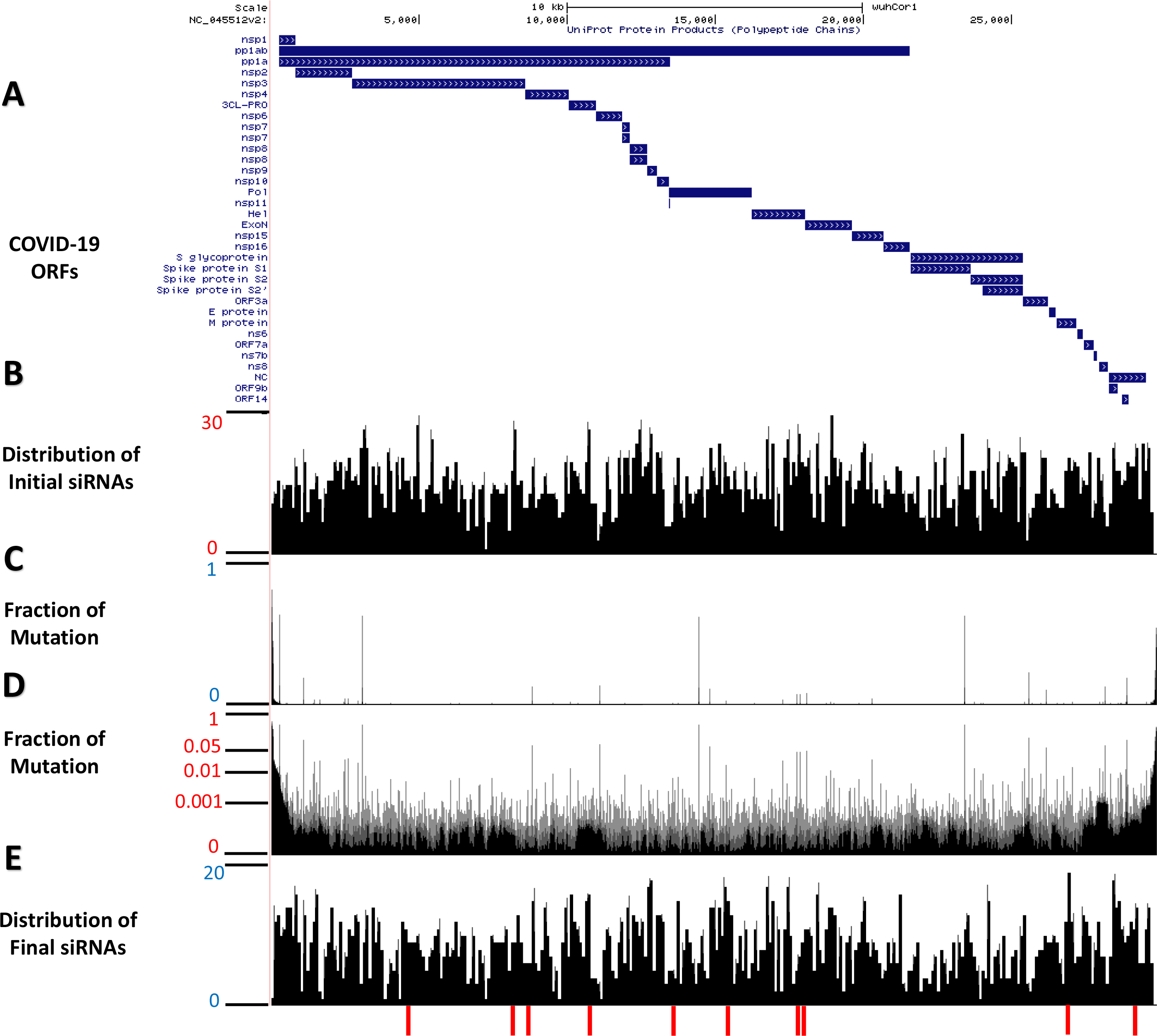
Distributions of designed siRNA against SARS-CoV-2. Mode A of the VirusSi pipeline was used to design siRNAs against SARS-CoV-2. **(A)** A schematic of the structure of the SARS-CoV-2 genome and predicted open-reading frames (ORFs). **(B)** A position-histogram of the distribution of initial siRNA candidates (after considering siRNA rules and SARS-CoV-2 folding structure) is shown. (**C, D**) The mutational profile of SARS-CoV-2 stains is shown with the vertical axis indicating the fraction of strains with mutation at the indicated position, plotted in (C) linear scale and (D) log scale. **(E)** A position-histogram of the distribution of the final siRNA candidates (after considering mutational profile and off-target effects) is shown. The bottom red bars indicate the positions of the top 10 candidate siRNAs.

To ensure that the candidate siRNAs could target most existing SARS-CoV-2 strains, we examined 15,920 SARS-CoV-2 genomes from the GISAID database to calculate the mutation profile. Based on the alignments between viral genomes, we computed the fraction of observed mutations (relative to the reference genome) for each base **(Figure 2C, 2D)**. The benefit of using a large number of SARS-CoV-2 genomes was revealed when we performed a similar analysis on 395 genomes from an earlier time of this pandemic. Substantial differences in genomic variation profiles were observed **(Supplementary Figure S1)**. After filtering out siRNAs that corresponded to more variable regions, we obtained 4432 siRNA candidates.

To reduce the potential side effects of candidate siRNAs on endogenous gene expression, we incorporated additional filtering and ranking in the VirusSi pipeline. Perfect base-pairing between endogenous genes and any siRNA can lead to undesired cleavage of endogenous transcripts, whereas matches between siRNA’s seed sequence with the endogenous transcript could lead to miRNA-like repressive effects. We first examined all candidate siRNAs to eliminate any that can perfectly match the human genome. We next penalized siRNAs based on their potential miRNA-like effects using the human lung transcriptome. Specifically, we evaluated all 8-mer binding sites (24) for a candidate siRNA in the 3’UTRs of human transcripts, and weighted them based on the level of expression of the transcripts in human lung tissue. These off-target penalties were further factored into a final siRNA score (see methods) for each siRNA candidate.

Based on the above steps, we came up with 2486 candidate siRNAs against SARS-CoV-2 (**Figure 2E**) with the location and sequences of the top ten candidates shown (**Figure 2E, Figure 3A**). Each of the top three siRNAs can target >99% of the 15,920 SARS-CoV-2 strains examined (**Figure 3B**), and a combination of four top siRNAs could ensure that each of the 15,920 SARS-CoV-2 strains had at least one siRNA targeting it **(Figure 3C**). The above steps were incorporated into the Mode A of the VirusSi pipeline (**Figure 1A**), which focuses on designing siRNAs against a single virus, with data inputs of the reference genome of the virus, genomes of different strains of the virus, as well as tissue transcriptomic data.

**Figure 3.**
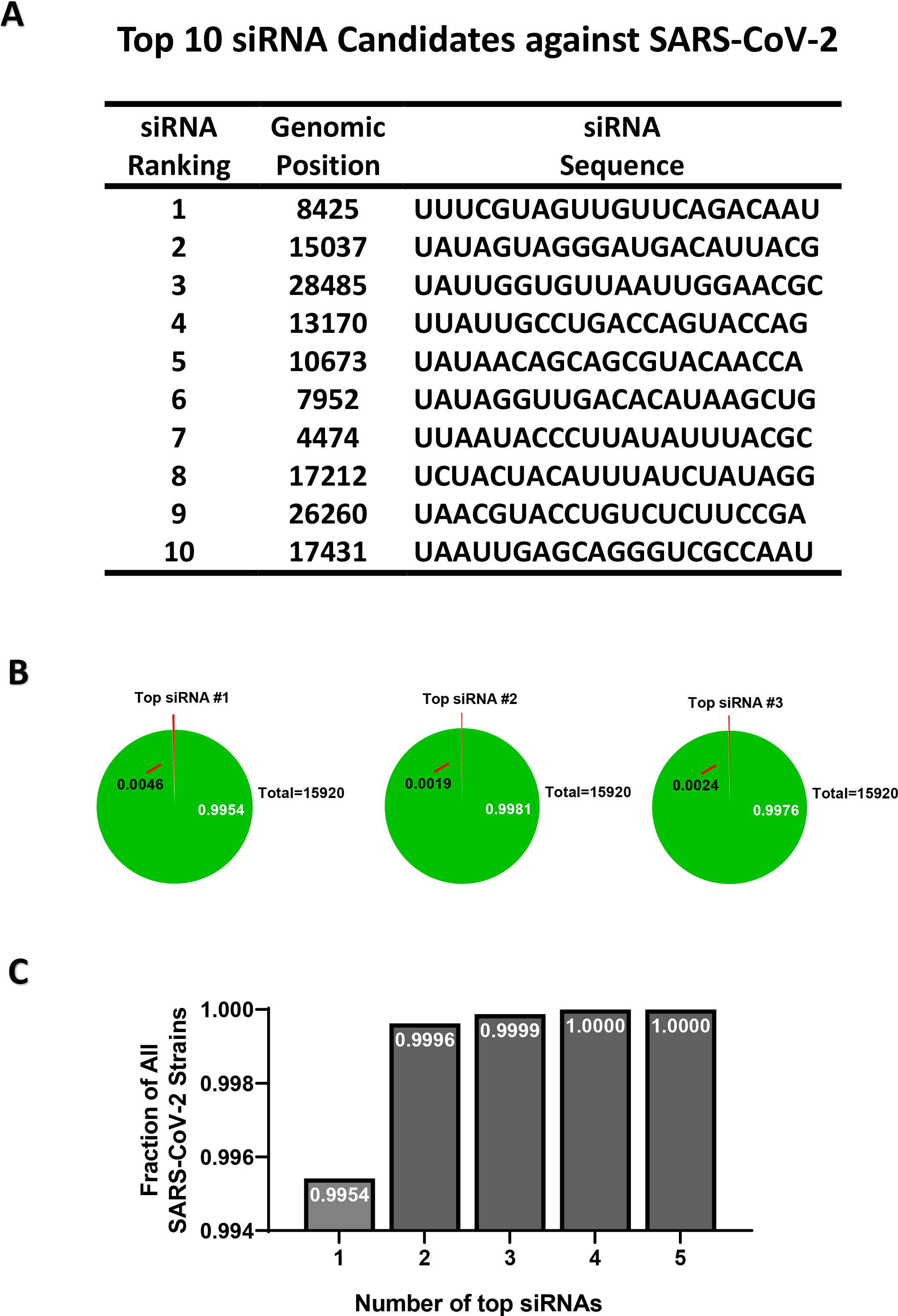
Properties of the top designed siRNAs against SARS-CoV-2. **(A)** The ranks, locations and sequences of the top 10 designed siRNAs are shown. **(B)** For each of the three top siRNAs, a pie chart is shown to indicate the fraction of SARS-CoV-2 strains that it could target (green area, white letters), or could not target (red area, arrow), with the total number of strains indicated as well. **(C)** The fractions of SARS-CoV-2 strains that could be targeted by at least one siRNA when considering a group of siRNAs all together. Groups plotted include only one siRNA (the top one), or top two, or up to the top 5.

In addition to designing siRNAs against SARS-CoV-2, we envisioned the possibility of establishing a reasonably small sized collection of pre-designed and safety-validated siRNA drugs that could be used off-the-shelf to target future respiratory viruses **(Figure 4A)**. This concept is rooted in the fact that any therapeutic siRNA drug would require pre-clinical and clinical trials for safety and efficacy that could take years and thus impractical to offer immediate response to any viral outbreak, if the siRNA design occurs after the outbreak. In contrast, pre-designed siRNAs could be pre-tested for safety and efficacy prior to the outbreak, and thus could be used off-the-shelf to combat a viral disease.

**Figure 4.**
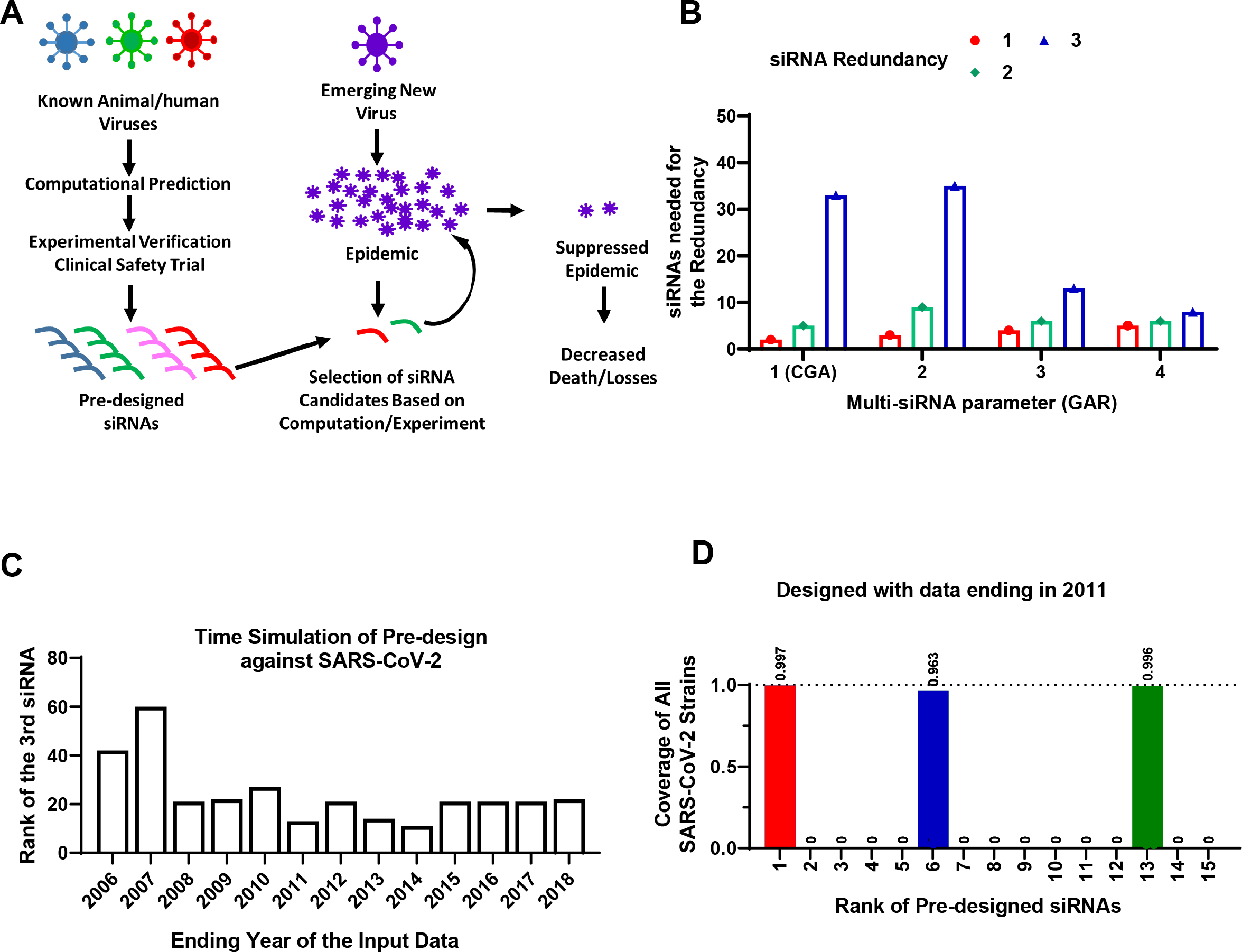
Pre-designing siRNAs against SARS-CoV-2. **(A)** A schematic of the model for pre-designing siRNAs to improve the speed of therapeutic response to future viral outbreaks. **(B)** Human and bat coronavirus genomes with an ending year of 2018 were used as input for the VirusSi pipeline, with the GAR algorithm and the Multi-siRNA parameters as indicated. When the Multi-siRNA parameter was 1, the results were identical to CGA. The numbers of top predicted siRNAs that were needed to identify one, two or three (redundancy) siRNAs against SARS-CoV-2 are plotted. (**C**) Time simulations were performed to evaluate the earliest time feasible to pre-design siRNAs against SARS-CoV-2. Human/bat Coronaviridae genomes with the indicated ending years were assembled and used as input for the VirusSi pipeline, using the SGAR algorithm. Data indicate the rank of the third siRNA that target SARS-CoV-2 (reflecting redundancy=3). **(D)** For the analysis from (C), the detailed data for input data ending in 2011 are plotted, with the top-ranked pre-designed siRNAs shown. Each bar represents the fraction of SARS-CoV-2 genomes that can be targeted by the corresponding siRNA, with the number above the bar indicating the exact fraction.

We thus expanded the VirusSi with a second mode, Mode B (**Figure 1B**), which takes the reference genomes from a class of existing viruses to predict a collection of siRNAs out of which some could target future viruses. Initially, we tested the seemingly straightforward strategy of identifying a collection of siRNAs, each of which could target the most numbers of input viral strains. However, using the SARS-CoV-2 outbreak as an example, none of the top 100 pre-designed siRNAs by this approach could target SARS-CoV-2 effectively (**Supplementary Table S2**, and see Methods). As an alternative, the Greedy algorithm (referred to hereafter as “conventional Greedy algorithm” or CGA), originally designed to solve the mathematical maximum coverage problem, could be used to identify the smallest number of siRNAs to ensure that each of the existing input viral strains could be targeted by at least one siRNA. By iterative evaluation of siRNAs to minimize the number of siRNAs to cover most strains (see flow chart in **Supplementary Figure S2A**), CGA could allow the possible prediction of at least one siRNA against a future virus, with the underlying assumption that any emerging future virus would bear some level of sequence similarity to one of the existing input viruses. However, for the purpose of building a collection of siRNAs against future viruses, having a single siRNA would not provide strong robustness in the system, because there could be failures of siRNA candidates due to issues of efficacy or safety(14). We thus developed the Greedy Algorithm with Redundancy (GAR) which reduced the penalty for siRNAs against viral strains that were targeted by already-selected siRNAs (see flow chart in **Supplementary Figure S2B**). By incorporating a Multi-siRNA parameter (see methods for details), GAR could thus tune the robustness of the prediction.

To evaluate both the feasibility of predicting siRNAs against future viruses, and to evaluate the relative performance of GAR versus CGA, we simulated the emergence of SARS-CoV-2. We downloaded Coronaviridae genomic sequences from the Virus Pathogen Database (25). These sequences were filtered so that only sequences deposited up to the end of 2018 were kept, a time point about a year prior to the COVID-19 outbreak. We further filtered to retain Coronaviridae strains that could use human or bat as a host, resulting in 717 coronavirus sequences. The inclusion of bat coronaviruses was due to bat as a natural reservoir of many coronaviruses that eventually crossed the species barrier into human (26). Using these sequences as input, which did not include any SARS-CoV-2 stains, we predicted and ranked the siRNAs using both CGA and GAR. We evaluated the performance of these two algorithms by counting the number of top predicted siRNAs that were needed to have at least one (redundancy=1), two (redundancy=2) or three (redundancy=3) non-overlapping siRNAs, each of which could target the majority of SARS-CoV-2 strains. Interesting, both GAR and CGA successfully predicted siRNAs that could target SARS-CoV-2 (**Figure 4B**). Comparing results from GAR and CGA (note that CGA is equivalent to running GAR with the Multi-siRNA setting of one), however, showed that with higher redundancy requirements (e.g. redundancy of 3), GAR performed substantially better than CGA with higher Multi-siRNA settings, at a minor cost of slightly worse performance for the redundancy requirement of one (**Figure 4B**). The analyses above used time simulation and demonstrated the feasibility of designing siRNAs against a future virus using existing viral information. The results further demonstrated that GAR outperformed CGA on the robustness of finding multiple siRNAs against a future virus.

The existing viral genome information tends to be heavily biased toward virus types that have been more intensely studied. On the other hand, it is not guaranteed that a future respiratory virus would emerge with strong similarity to an intensely studied virus in the past. Both GAR and CGA treated each input stain equally, without considering similarity among input stains, and thus could be more susceptible to the biases in existing data. Hence, we reasoned that developing SGAR, or Similarity-weighted Greedy Algorithm with Redundancy, could be potentially beneficial to mitigate the effects of such biases. We designed SGAR to add weights based on similarity between input viral strains, with strains that differ more from other input sequences weighted more (see flow chart in **Supplementary Figure S2C**). To simulate biased input data, we randomly selected a series of 100 input Cornoaviridae viral strains out of the Virus Pathogen Database, so that (1) different ratios of human/bat coronaviruses were added, because such sequences improved prediction results for siRNAs against SARS-CoV-2 using GAR (**Supplementary Figure S3**), and (2) other than human/bat Coronaviridae sequences, no other beta-coronaviruses were included. Each of the 100-input-strain sets was subjected to GAR and SGAR predictions. Comparing results between GAR and SGAR, significantly fewer siRNAs were needed for SGAR to identify SARS-CoV-2-targeting siRNAs when lower percentage of human/bat coronavirus sequences were included, whereas GAR’s performance trended a little better, albeit not statistically significant, when all input sequences were human/bat viruses (**Supplementary Figure S4**). We thus included both GAR and SGAR in the VirusSi pipeline, which could be selected by the user.

We next asked when was the earliest time possible to pre-design siRNAs against SARS-CoV-2 by simulating the time cutoff for input viral strains. We again took the Coronaviridae sequences as input for SGAR, but limited the input sequences to an ending year that range from 2003 to 2018. We used an evaluation criterion of at least three siRNAs against SARS-CoV-2, to tolerate potential failures in efficacy and safety trials. We found that for the ending year from 2008 to 2018, a maximum of 27 top ranked siRNAs was needed, whereas predictions using data ending before 2008 had worse performances (**Figure 4C**). As an example, with data ending in year 2011, only 13 siRNAs were needed to find three siRNAs, each of which was predicted to target >96% of SARS-CoV-2 strains (**Figure 4C, 4D, Supplementary Table S3**). In hindsight, if this pipeline were available in 2009 or before, there could have been at least a 10-year lead period to prepare against the SARS-CoV-2 outbreak. These data suggest that the pre-designing concept holds a strong promise.

We next asked whether we could predict siRNAs against MERS-CoV and SARS-CoV-1, using other coronavirus strains. These two viruses were the pathogens causing MERS and the SARS epidemics, which started in June 2012 and November 2002, respectively. We first used Coronaviridae sequences prior to 2019, but removed MERS-CoV or SARS-CoV-1 sequences. When using SGAR on these two input sequence sets, 8 and 19 top siRNAs were needed to identify three siRNAs that could target MERS-CoV and SARS-CoV-1, respectively (**Supplementary Figure S5A, S5B**). We next performed time-simulation of pre-designing siRNAs against MERS-CoV. We used human and bat coronaviruses ending in 2011, with a total of 235 strains, which were noticeably fewer than the number for 2018. Using these input stains, the first three siRNAs against MERS-CoV were ranked 12, 26 and 52 among all predicted siRNAs, each of which could target all MERS-CoV stains (**Supplementary Figure S6**). This meant that 26 siRNAs were needed to identify two MERS-CoV-targeting siRNAs, Predictions using data ending in year 2010 or before had weaker performances (**Supplementary Table S4**). Nevertheless, two MERS-CoV-targeting siRNAs could be identified among the top 100 predicted siRNAs using data ending as early as 2008. The reduced performance was associated with reduced number of input strains, with 218 stains for data ending in 2010 and 163 strains for data ending in 2008. We next tested pre-designing against SARS-CoV-1. Only 54 human/bat coronavirus sequences were available prior to 2002. Among the Top 100 SGAR-predicted siRNAs based on such input sequences, none could target SARS-CoV-1. These data indicate that pre-designing siRNAs against MERS-CoV can be achieved using sequence information prior to the outbreak. At the same time, out data suggest that the scarcity of human/bat coronavirus sequences in early years weaken the performance of the algorithm and makes it difficult to pre-design siRNAs against SARS-CoV-1.

We next asked whether pre-design is feasible against a respiratory virus that is not a coronavirus. We thus explored pre-design against the 2009-H1N1 influenza virus, which underlies the 2009 swine flu pandemic. We took type-A influenza genomes as inputs, with the input-data ending year ranging from 2000 to 2008. To make the pre-design more challenging, we removed all other H1N1 strain sequences from the input. Results show that pre-design was possible even with input data ending in year 2000 (1093 total input strains), when each of the top three pre-designed siRNAs was capable of targeting all 2009-H1N1 strains (**Supplementary Table S5**).

Some pre-designed siRNAs could fail pre-clinical experimental testing, which would lead to fewer siRNAs entering clinical trials and thus reduce the robustness to prepare for future outbreaks. To overcome this problem, we added a functionality in the VirusSi pipeline to redesign siRNAs based on prior experimental evaluation status. This was achieved by having two additional inputs for the pipeline, namely a list of “successful siRNAs” and a list of “failed siRNAs”. We simulated the re-design process using SARS-CoV-2 as an example. We used the pre-designed siRNAs against SARS-CoV-2, with data ending in 2018, which is the same as the input data used in **Figure 4B**. Among the top 25 pre-designed siRNAs, there were four siRNAs that could target > 94% of SARS-CoV-2 strains. We randomly assigned these 25 siRNAs into the successful and failed groups, with a ~ 50% success rate, leading to three out of four SARS-CoV-2-targeting siRNAs falling into the failed group. Redesigning through the VirusSi pipeline still yielded four siRNAs that could target > 94% of SARS-CoV-2 strains among the top 25 siRNAs, including the one that was labeled as successful (**Supplementary Table S6**). These data support that additional siRNAs could be easily re-designed to supplement failed siRNAs, to facilitate preparation against future viruses.

Lastly, we used SGAR to pre-design siRNAs against potential future coronaviruses using currently existing coronavirus sequences. We first used human/bat coronaviruses as an input. We combined 769 non-SARS-CoV-2 human/bat coronavirus strains and 192 SARS-CoV-2 strains (20% of total input strains) as input data, and run VirusSi with a Multi-siRNA setting of 3. The top 30 and 100 pre-designed siRNAs are shown in **Figure 5A** and **Supplementary Table S7**, respectively. Of note, among the top 16 siRNAs within this list, there were three that target the majority of SARS-CoV-2 stains (**Figure 5B**). In addition to the use of human/bat coronaviruses, we also pre-designed siRNAs using all Coronaviridae sequences (with 800 subsampled SARS-CoV-2 strains), totally 4000 strains. The top 100 predicted siRNAs are listed in **Supplementary Table S8**. Interestingly, out of the top 30 pre-designed siRNAs based on human/bat coronaviruses alone, there were 20 that overlapped with the second list based on all coronaviruses (**Supplementary Table S7**). We propose that these two lists of siRNAs could form candidates to prepare for future coronavirus outbreaks.

**Figure 5.**
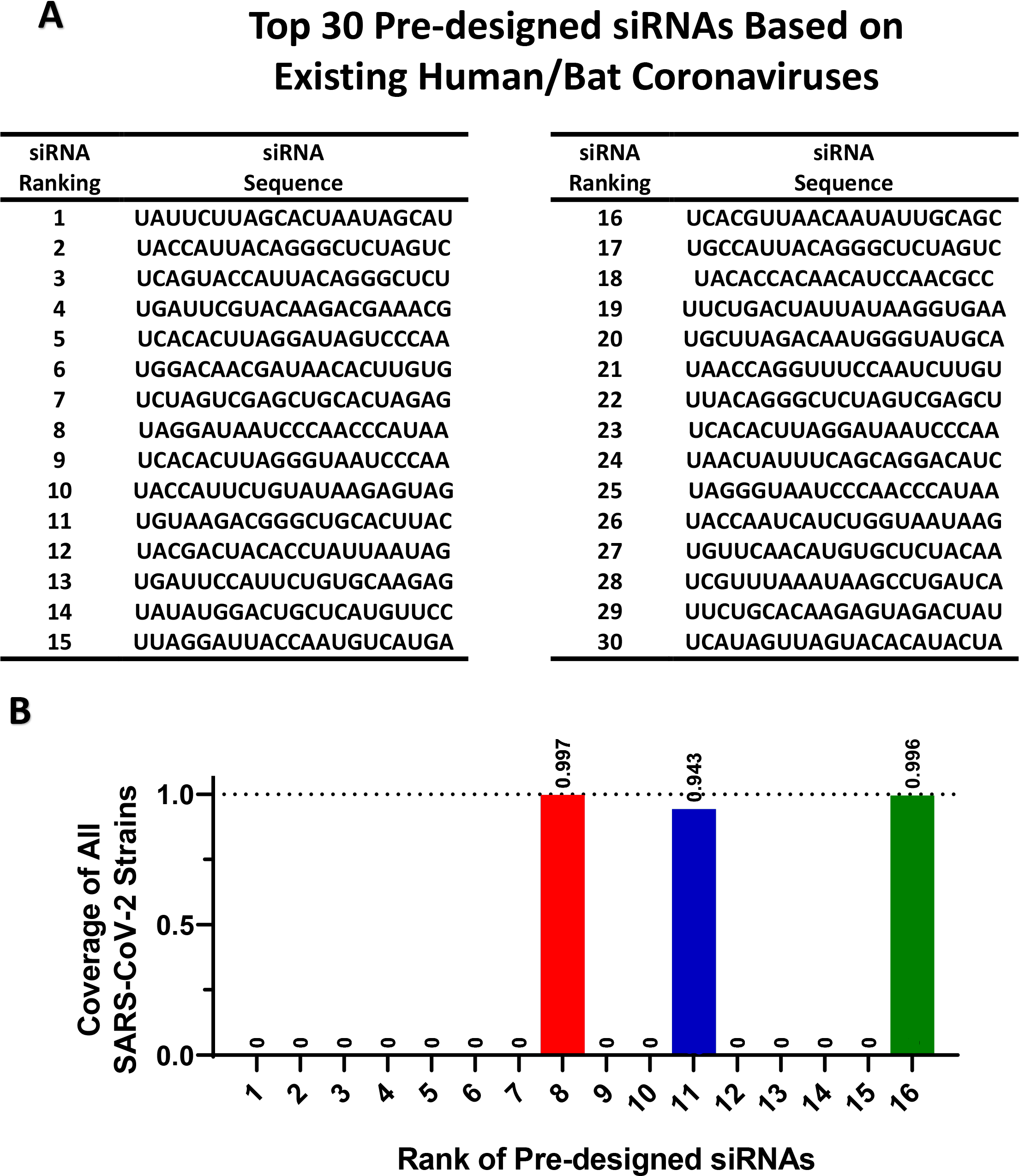
Pre-designing siRNAs against future coronavirus based on existing Coronaviridae genomes. **(A)** The existing human/bat Coronaviridae genomes were used as input for the VirusSi pipeline using the SGAR algorithm with a Multi-siRNA setting of 3. The top 30 predicted siRNAs are shown. **(B)** For the analysis from (A), the siRNAs were examined for their ability to target SARS-CoV-2. Each bar represents the fraction of SARS-CoV-2 genomes that can be targeted by the corresponding siRNA, with the number above the bar indicating the exact fraction.

## Discussion

In this study, we developed the VirusSi pipeline and used time simulation to demonstrate the feasibility of using VirusSi to pre-design siRNAs against future respiratory coronaviruses. VirusSi can also rationally design siRNA against an existing virus. The prospect of pre-designing anti-viral siRNA reagents is exciting, which has not been explored previously. We demonstrated the pre-design of siRNAs against SARS-CoV-2, MERS-CoV and 2009-H1N1 using data prior to their outbreaks. In the case of SARS-CoV-2, we showed the feasibility of pre-designing using data with a cutoff as early as 2008. For SARS-CoV-1, the pre-design using data prior to 2002 did not generate successful siRNAs. We believe the reason is the limited number of human/bat coronavirus genomes prior to 2002, because including later strains (without SARS-CoV-1 sequences) made it possible to design siRNAs against SARS-CoV-1. Similar observations were made for MERS-CoV. Although pre-design of siRNAs against MERS-CoV was possible, performance was substantially improved when later coronavirus strains (without MERS-CoV sequences) were included. These data also strongly argue for the need to sustain and enhance the basic research into respiratory viruses in animal species including bats. Our data suggest that the more information is available, the better the pre-design concept can work.

We envision a model in which pre-designing could allow testing for efficacy and safety prior to future outbreaks of newly emerged viruses, and thus accelerating responses toward such viruses. Our data of pre-design against SARS-CoV-2, MERS-CoV and 2009-H1N1 suggest that a collection of a small number (~30) of pre-designed siRNAs could have a good likelihood of targeting future viruses. The small number is key, because clinical trials will be both expensive and time consuming. To make use of the pre-designed siRNAs, efforts first need to be put into validating the efficacy of each siRNA, possibly using synthetic targets as a readout through *in vitro* experiments in cell culture, as well as using existing viral strains if they match the designed siRNA. This should be a reasonably fast process that can determine which pre-designed siRNAs should be weeded out due to the lack of targeting efficacy. As we demonstrated, additional siRNAs could be quickly designed using VirusSi to replace such failed siRNAs. siRNAs that successfully met efficacy criteria could undergo further testing using animal models. Eventually, siRNAs could be tested in human clinical trials, at least for safety.

As mentioned above, the designed siRNAs in this study will need to be tested in a wet-lab setting for both efficacy and safety. RNA secondary structure, which is difficult to predict accurately across long RNA sequences, and the presence of RNA binding proteins, may affect siRNA efficacy. Similarly, while our pipeline takes consideration of predicted off-target effects, the full spectrum of off-target effects could only be revealed in *in vivo* experimentation, possibly through both pre-clinical studies and clinical trials. Additionally, while the delivery of siRNAs into lung cells *in vivo* has been demonstrated in animal models (9–12) and has been shown to reduce virulence in two SARS-CoV-2 models *in vivo* (14), delivery of multiple siRNAs into lung cells with high delivery efficiency still require intense experimentation before such siRNAs could be used as anti-viral therapeutics. Both SARS-CoV-1 and SARS-CoV-2 harbor gene products that have activities to inhibit the siRNA machinery (27, 28). Encouragingly, it has been shown that siRNAs against SARS-CoV-1 and SARS-CoV-2 can be effective in experimental settings (14, 29–31).

The Mode B of the VirusSi pipeline utilizes GAR and SGAR algorithms developed in this study. Both GAR and SGAR have a Multi-siRNA setting to improve the robustness of identifying more than one siRNAs against a future viral strain within a small number of topranked siRNAs. Our data show that SGAR have some advantage over GAR suggesting that SGAR is the more preferred mode. Nevertheless, both GAR and SGAR are included in the VirusSi pipeline, and the choice of the algorithm as well as the choice of the Multi-siRNA setting could be defined by users. Lastly, while the VirusSi pipeline was developed for designing siRNAs and used in this study against respiratory viruses, minor modification of the pipeline could permit the design of crRNAs for the Cas13d system, which has been demonstrated to both detect virus and act as anti-viral tools (7). Designing siRNAs against non-respiratory viruses, such as the EBOLA virus (8), should also be possible using VirusSi.

## Methods

### Source of Genomic Data

Genome data, their annotation, and the data download dates are listed below.

| Data Type | Data Source | Download Date<br>(year.month.day) |
| --- | --- | --- |
| SARS-CoV-2 genomic<br>sequence data | GISAID Database | 2020.5.4 |
| SARS-CoV-2 reference genome<br>sequence data (Accession<br>Number: NC_045512) | NCBI GenBank | 2020.5.4 |
| Genomic sequence data of<br>Coronaviridae | Virus Pathogen Database<br>and Analysis Resource | 2020.6.18 |
| Gene annotation | GENCODE (v34) | 2020.5.5 |
| Gene expression data for human<br>lung | GTEx(v8) | 2020.5.22 |
| Influenza genomic sequence<br>data | Influenza Virus Database | 2020.8.12 |

### Mode A pipeline of VirusSi

The mode A of the VirusSi pipeline focuses on designing siRNA candidates against a single virus, utilizing genomic sequences of multiple strains of the virus as an input. The methods below use SARS-CoV-2 as an example, but the pipeline can be used for any RNA virus.

#### 1. Rules for siRNA candidates

The program scanned with a 21nt siRNA target window along the viral RNA genome. For SARS-CoV-2, the scan started at 25 nt from the 5’ end and stopped 130 nt from the 3’ end, to avoid low quality regions in the genomes. For Influenza, the scan started at 25 nt from the 5’ end and stopped 25 nt from the 3’ end of each RNA segment. The program had two sets of rules that the user could choose from. For the first set of rules (default setting, modified from GPP Web Portal), target sequence would be excluded from further consideration if it met any of the following criteria: (1) the identity of the 3’ nucleotide is not an A; (2) the identity of the 5’ di-nucleotides is AA; (3) the sequence contains a run of four of the same nucleotide in a row; (4) sequence has non-optimal GC content (GC <= 25% or > 60%); (5) sequence contains ambiguous bases (e.g. N); (6) sequence has a run of 7 consecutive C/G nucleotides; (7) unpaired probabilities of the last 8bp of candidate targeting region is larger than 0.01157 (based on RNAxs(24)). For each target sequence that was not excluded based on the above criteria, a score was calculated based on rules in Supplementary Table S1.

The second rule set used the exact rules in RNAxs(24). The siRNA candidate criteria include: (1) the accessibility probability on the last 8nt of the targeted region is larger than or equal to 0.01157; (2) the accessibility probability on the last 16nt of the targeted region is larger than or equal to 0.001002; (3) the energy asymmetry is larger than or equal to 0.4625; (4) the sequence asymmetry is larger than or equal to 0.5; (5) the self-folding energy is larger than or equal to 0.9; (6) the free end parameter is larger than or equal to 0.625. Users could select rule sets in the pipeline and the first rule set is the default rule set which was used in this study.

#### 2. Filtering and penalizing candidate siRNAs based on mutational profiles of SARS-CoV-2

We first used the reference genome of SARS-CoV-2 from NCBI GenBank and aligned the genomic sequence of each of the downloaded 15,920 COVID-19 strains with it, using the GISAID database (32). Based on the alignment result, we documented all mutations present in each strain. Both indel and point mutations were included in the mutation tally. For some sequences, there were ambiguous bases (N and other non-ACGT bases). We considered such bases as unreliable. Mutational rate was calculated as the ratio of the number of mutations occurring at a given genomic base to the total number of strains for which genomic data were reliable at that base. Mutational profile consisted of mutational rates for each of the base in the reference SARS-CoV-2 genome.

We excluded candidate siRNAs whose targeting region had an overall mutation rate of more than a threshold of 0.05. The overall mutation rate for a given siRNA was calculated as follows: the number of SARS-CoV-2 strains which had at least one mutation in the targeting region and whose sequence was reliable in the targeting region, divided by the number of SARS-CoV-2 strains whose sequences were reliable in the targeting region.

After that step, the mutation-related siRNA score was calculated as (0.05 - overall mutation rate) / 0.05. Then we ranked the siRNAs based on the mutation-related siRNA scores in a descending order. We also assigned a mutation-related rank value to each siRNA based on the order. If there was a tie between multiple siRNAs with the same mutation-related siRNA scores, they received the same rank.

#### 3. Filtering and Penalizing candidate siRNAs with predicted strong off-target effects

We considered the expression levels of transcripts in human lung tissues to determine the off-target score. Lung tissue expression data were downloaded from GTEx database, with a total of 867 lung samples. Host transcripts with a mean expression level lower than 0.05 (TPM) were not be considered in this step. For the rest of the transcripts, they were ranked by mean expression levels in human lung tissue and were binned into five equal-sized bins, and assigned an expression-level adjustment factor from 1 to 5 with higher expression bins having higher adjustment factors.

We considered a gene to be potentially off-target if it had one of more pairing regions for a siRNA. There were three types of pairings considered in the off-target analysis: perfect match sites, imperfect match sites and 8mer match sites. First, any candidate siRNA with a perfect match site in the human transcriptome (as defined by GENCODE) was eliminated. The second type was imperfect match sites for a siRNA, which was defined as a putative binding site of the siRNA with a perfect match involving the first 8 bases of the siRNA and up to three unmatched bases in the rest of the siRNA. Each gene having imperfect match site was multiplied by the corresponding expression-level adjustment factor, and their sum became the final imperfect match score. We excluded a candidate siRNA if it had an imperfect match score of more than 20. If the number of imperfect match score was larger than 0 but less than 20, the imperfect match adjustment factor was calculated as: (20 - imperfect match score) / 20. The third type was 8mer miRNA-like sites, which were defined according to Friedman *et al*. (24). Each 8mer site was multiplied by the corresponding expression-level adjustment factor, and the sum of them became the final 8mer match score. We excluded a siRNA when it had an 8mer match score of more than 20,000. If the 8mer match score was larger than 0 but less than 20,000, we defined the 8mer adjustment factor to be (20000 – 8mer match score) / 20000.

After that step, the off-target-related siRNA score was calculated as the multiplication product of the imperfect-match adjustment factor and the 8mer adjustment factor. Then we ranked the siRNAs based on the off-target-related siRNA score for each siRNA in a descending order. We also assigned an off-target-related rank value to each siRNA based on the order. If there was a tie between multiple siRNAs with the same off-target-related siRNA scores, they received the same rank.

#### 4. Final siRNA score and the ranking of siRNA candidates

After these three steps, the final siRNA ranking was calculated based on the three different rank values as stated above. The siRNAs were then ranked based on the maximal value of the three individual rank values in ascending order. If there were siRNAs with a tie of the maximal rank, the average rank of the three individual ranks was used as a tiebreaker. The final rank was used as the final score for the siRNA.

### Mode B pipeline of VirusSi

The mode B of the VirusSi pipeline focuses on pre-designing siRNA candidates against future emerging viruses, with the input being the genomic sequences of a class of viruses. The methods below use coronavirus as an example, but the pipeline could be used for other classes of RNA viruses.

#### 1. Input viral genome information

We combined the genome sequences for input viral strains (details described in the sections below) in a single fasta files and used this file as the input of the pipeline.

#### 2. Designing candidate siRNAs for each of the input viral genome

a. **Rules for siRNA candidates** Rules used as the same as in the Mode A pipeline.
b. **Filtering and penalizing candidate siRNAs with predicted strong off-target effects** Filtering and penalizing for off-target effects were the same as in Mode A. All the siRNAs which did not meet the requirement of off-target cutoff were not considered in Mode B.

#### 3. Using the GAR and SGAR algorithm to rank the siRNAs based on the input viral genomes

a. **SGAR** The SGAR algorithm has the following steps. Step 1: Calculate similarity between input strains using overlaps of candidate siRNAs. We calculated the similarity between viral strains by quantifying the overlap of candidate siRNA between any two input strains:

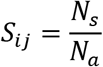

*S_ij_*: siRNA target similarity between viral stains *i* and *j*.
*N_s_*: the number of siRNAs shared by the two stains *i* and *j*.
*N_a_*: the union of all unique siRNAs targeting the two stains; Step 2: Calculate a similarity weighted strain score, for each input strain

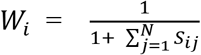

*W_i_*: Weight for stain i
*S_ij_*: similarity between stains *i* and *j*.
*N*: Total number of input strains Step 3: Start the greedy selection Obtain the union of all siRNAs targeting any of the input strains. For each siRNA, the target score was calculated as:

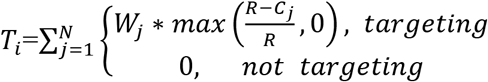

*T_i_* : Target score for siRNA #i
*W_j_*: The weight of strain j
*R*: The user-defined Multi-siRNA parameter
*C_j_* : Count of selected siRNAs that can target strain j.
*N*: Total number of input stains Sort all the siRNAs based on the target score, in a descending order. The first siRNA selected was the one that had the top target score. If multiple siRNAs had the same top score, one of them was randomly picked. Step 4: Iterative selection of additional siRNAs For each of the remaining siRNAs, re-calculate the target score, utilizing the information on which strains could be targeted by the already selected siRNAs. The siRNA with the highest targeting score was selected as the second selected siRNA. This process reiterated, until the number of selected siRNAs reached a pre-defined number of output siRNAs, which was set to 100 in this study.
b. **GAR** GAR is similar to the SGAR algorithm, but there was no weighting for the stains based on similarity. The target score for a siRNA was defined as:

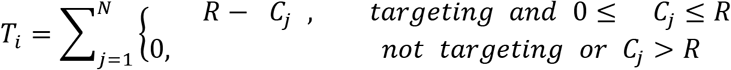

*T_i_* : The target score for siRNA #i;
*R*: The user-defined Multi-siRNA parameter
*C_j_* : Count of selected siRNAs that can target strain j.
*N*: Total number of input stains
c. **CGA (Conventional Greedy Algorithm)** CGA was implemented by setting the Multi-siRNA parameter R to be 1 in the GAR algorithm.

### Evaluating GAR, SGAR and CGA, and using SGAR to pre-design siRNAs

#### 1. Evaluation of GAR versus CGA

To compare the performance of GAR vs CGA, we performed stimulations. We used data from the Virus Pathogen Resource (ViPR) database as input information for prediction, and used 15,920 SARS-CoV-2 data from GISAID database for the evaluation of the prediction results. The input data were Coronaviridae genomes collected before or in 2018. We further confirmed the removal of the bat coronavirus sequence RaTG13 (GenBank MN996532) due to its later sequence deposit. VirusSi pipeline was run with the GAR algorithms, using the Multi-siRNA parameters ranging from 1 to 4. When the Multi-siRNA parameter was 1, the algorithm and results were identical to CGA.

After the prediction, we counted siRNAs which could target SARS-CoV-2 strains. An siRNA was considered SARS-CoV-2-targeting if it could perfectly match > 50% of SARS-CoV-2 strains. Additionally, to avoid multiple siRNAs with overlap in their target site locations, and hence not independent from each other, we removed any siRNA that overlapped with a better ranked siRNA.

#### 2. Evaluation of SGAR vs GAR

Evaluation of SGAR vs GAR was performed similar to the comparison of GAR vs CGA, with the following differences. The first difference was the input data. To build input data, we divided all Coronaviridae genome data into two data types and mixed them in different proportions. The first data type included genome sequences of human and bat viruses in Coronaviridae. The second data type included genome sequences of all non-human/bat viruses in Coronaviridae, with genomes of betaCoVs removed to enhance strain biases. Type 1 input data and Type 2 input data were mixed, so that the proportion of Type 2 sequences was 0%, 10%, 20%, 40%, 60%, 80% or 100%. For each mixing ratio, we generated 5 randomly chosen sequence sets of 100 strains per set. Each input sequence set was then subjected to the VirusSi pipeline, using SGAR and GAR, in separate runs. We used the default Multi-siRNA setting of 3.

To compare results between GAR and SGAR, we performed Wilcoxon matched-pairs signed rank test, using GraphPad Prism 8 software. Since in this analysis we limited the number of output siRNAs to be 100, if none of the top 100 siRNAs could target SARS-CoV-2, we set 100 as the number for the sake of statistical comparison.

#### 3. Time simulation of pre-designing siRNAs against SARS-CoV-2, MERS-CoV, SARS-CoV-1, and 2009-H1N1

For the time simulation of pre-designing against SARS-CoV-2, MERS-CoV and SARS-CoV-1, we assembled multiple sets of input data using all Coronaviridae genomes from the ViPR database, with an ending collection time from year 2002 to 2018. We then used the VirusSi pipeline, with the SGAR algorithm, to predict siRNAs using each of the input datasets. We used the default Multi-siRNA setting of 3. Evaluation of performance on SARS-CoV-2-targeting siRNAs was carried out in the same way as described in the section comparing GAR vs CGA. For evaluating results on MERS-CoV, we used 251 MERS-CoV genomes from the ViPR database. For evaluating results on SARS-CoV-1, we used 52 SARS-CoV-1 genomes from the ViPR database.

For the time simulation of pre-designing against 2009-H1N1, we assembled multiple sets of input data using all type-A Influenza genomes from the Influenza Virus Database, with an ending collection time ranging from year 2000 to 2008. We further removed any H1N1 strains from these collections. We then used the VirusSi pipeline, with the SGAR algorithm, to predict siRNAs using each of the input datasets. We used the default Multi-siRNA setting of 3. Evaluation of performance on 2009-H1N1 was similar to that of SARS-CoV-2, but used a total of 23 2009-H1N1 genomes from the NCBI GenBank.

#### 4. Pre-designing siRNAs against future coronavirus based on existing coronavirus strains

Coronaviridae genomes from the ViPR database were first divided into those of SARS-CoV-2 and non-SARS-CoV-2 strains. For the first pre-design, we combined 769 human/bat non-SARS-CoV-2-strains with 192 SARS-CoV-2 strains, with the latter corresponding to 25% of the non-SARS-CoV2-2 strains. The reason for only including a fraction of SARS-CoV-2 genomes was due to the very large number of SARS-CoV-2 strains available, which would likely distort the pre-design results. The VirusSi pipeline was run with SGAR, with the default Multi-siRNA setting of 3.

For the second pre-design, we used 3200 non-SARS-CoV-2 Coronaviridae genomes available in the database, regardless of the host. These genomes were combined with 800 SARS-CoV-2 genomes (representing 25% of the non-SARS-CoV-2 genomes) as input. The VirusSi pipeline was run with SGAR, with the default Multi-siRNA setting of 3.

#### 5. Simulating the re-design of siRNAs based on experimental results

We first performed pre-design against SARS-CoV-2, using human/bat Coronaviridae genomes with an ending year in 2018, as described in the time-simulation section above. For the resulting 25 top pre-designed siRNAs, we randomly labeled 13/25 (~50%) to be successful, with the rest being failed. We re-designed using VirusSi, by inputting the lists of successful and failed siRNAs. The re-design process was similar to the pre-design procedures, with the following differences. First, any selected siRNA that overlapped with the successful list was automatically retained. Second, for any selected siRNA that was a member of the failed list, this siRNA was replaced with another siRNA based on the following. (1) The replacement siRNA was not a member of the successful list or failed list. (2) The replacement siRNA was not an siRNA already selected in the SGAR process. (3) The replacement siRNA had the highest target similarity score with the siRNA to be replaced. The target similarity score was calculated based on strains that could be targeted by both the replacement siRNA and the siRNA to be replaced (referred to as “shared target”), and was defined as:

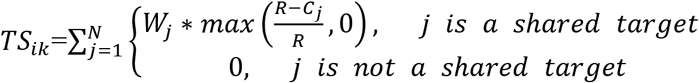

*TS_ik_*: Target similarity score between siRNA #i and #k
*W_j_*: The weight of strain j (see section on SGAR)
*R*: The user-defined Multi-siRNA parameter (see section on SGAR)
*C_j_*: Count of selected siRNAs that can target strain j. (see section on SGAR)
*N*: Total number of input stains (see section on SGAR)

### Pre-designing siRNAs based on broadly targeting siRNAs

In addition to CGA, GAR and SGAR, we also performed tests to pre-design siRNAs based on selecting siRNAs that had the broadest targeting capability on input viral sequences. For this analysis, the input strains were the same as used in the section comparing GAR vs CGA. The input data were subjected to the Mode B of the VirusSi pipeline, but the ranking of the siRNAs was not based on CGA, GAR or SGAR. Instead, siRNAs were ranked based on the number of input strains they could target in a descending order. The top 100 siRNAs from this rank list was used to evaluate performance on SARS-CoV-2. Evaluation of performance was carried out in the same way as described in the section comparing GAR vs CGA.

### Availability of the computational pipeline

The VirusSi pipeline is available as a command-line software, which has been deposited in the GitHub (https://github.com/dingyaozhang/VirusSi).

## Acknowledgements

We thank the contributors to the GISAID database (**Supplementary Table S9**) and all databases that have been used in this study. We thank colleagues within Yale Cooperative Center of Excellence in Hematology for stimulating discussions and suggestions. J.L. was supported in part by NIH grants (R01GM116855, R33CA225863 and R33CA246711).

---

**Figure S1. Comparison of mutation profiles using different numbers of SARS-CoV-2 strains.** Available SARS-CoV-2 strains ending January 31st, 2020 (395 strains, top panel) and those ending May 4th, 2020 (15920 strains, bottom panel) were compared to the reference SARS-CoV-2 genome. The fractions of mutations at each genomic position are plotted. Red boxes indicate examples of strong differences between the two profiles.

**Figure S2. Flow charts of CGA, GAR and SGAR algorithms.** The flow charts indicating the detailed steps in the algorithm are shown.

**Figure S3. Human/bat coronavirus sequences improve pre-design of siRNAs against SARS-CoV-2.** Coronaviridae genomes with an ending year of 2018 were divided into those that were human/bat coronavirus or those that were not. Each input dataset was randomly drawn so that it contained 100 genomes and the indicated percentage of human/bat coronavirus genomes. Five input datasets were used for each mixing ratio. VirusSi pipeline was run with GAR and the Multi-siRNA setting of 3, with total of 100 output siRNAs. The numbers of top siRNAs that were needed to identify three SARS-CoV-2 targeting siRNAs are shown. If none of the top 100 siRNAs could target SARS-CoV-2, the number in the plot was set to 100. Error bars represent standard deviations.

**Figure S4. Comparison of GAR vs SGAR using biased input data.** Coronaviridae genomes with an ending year of 2018 were divided into those that were human/bat coronavirus or those that were not. BetaCoVs were further removed from the non-humna/bat virus set to increase bias. Each input dataset was randomly drawn so that it contained 100 genomes and the indicated percentage of human/bat coronavirus genomes. Five input datasets were used for each mixing ratio. VirusSi pipeline was run with GAR and the Multi-siRNA setting of 3, with total of 100 output siRNAs. The numbers of top siRNAs that were needed to identify two SARS-CoV-2 targeting siRNAs were quantified, and the difference in the rank of the 2^nd^ siRNA between GAR and SGAR algorithms are shown in a whisker plot. Significance comparing GAR vs SGAR across the indicated groups (bar on the top) was calculated using the Wilcoxon matched-pairs signed rank test. *: p< 0.05.

**Figure S5. Predicting siRNAs against MERS-CoV and SARS-CoV-1.** SARS-CoV-1 (for A) or MERS-CoV (for B) sequences were removed from Coronaviridae genomes with an ending year of 2018. The VirusSi pipeline with the SGAR algorithm was run on these input data, using a Multi-siRNA setting of 3. **(A)** Top ranked siRNAs were evaluated for their ability to target SARS-CoV-1. Each bar represents the fraction of SARS-CoV-1 genomes that can be targeted by the corresponding siRNA, with the number above the bar indicating the exact fraction. **(B)** Similar data as in (A) are shown for MERS-CoV.

**Figure S6. Pre-designing siRNAs against MERS-CoV.** Time simulations were performed to evaluate the earliest time feasible to pre-design siRNAs against MERS-CoV. Coronaviridae genomes with 2011 as the ending year were assembled and used as input for the VirusSi pipeline, using the SGAR algorithm with a Multi-siRNA setting of 3. Top ranked siRNAs were evaluated for their ability to target MERS-CoV. Each bar represents the fraction of MERS-CoV genomes that can be targeted by the corresponding siRNA, with the number above the bar indicating the exact fraction.

