## Supplementary Figures 1-6 for "*In Silico* Design of siRNAs Targeting Existing and Future Respiratory Viruses with VirusSi"

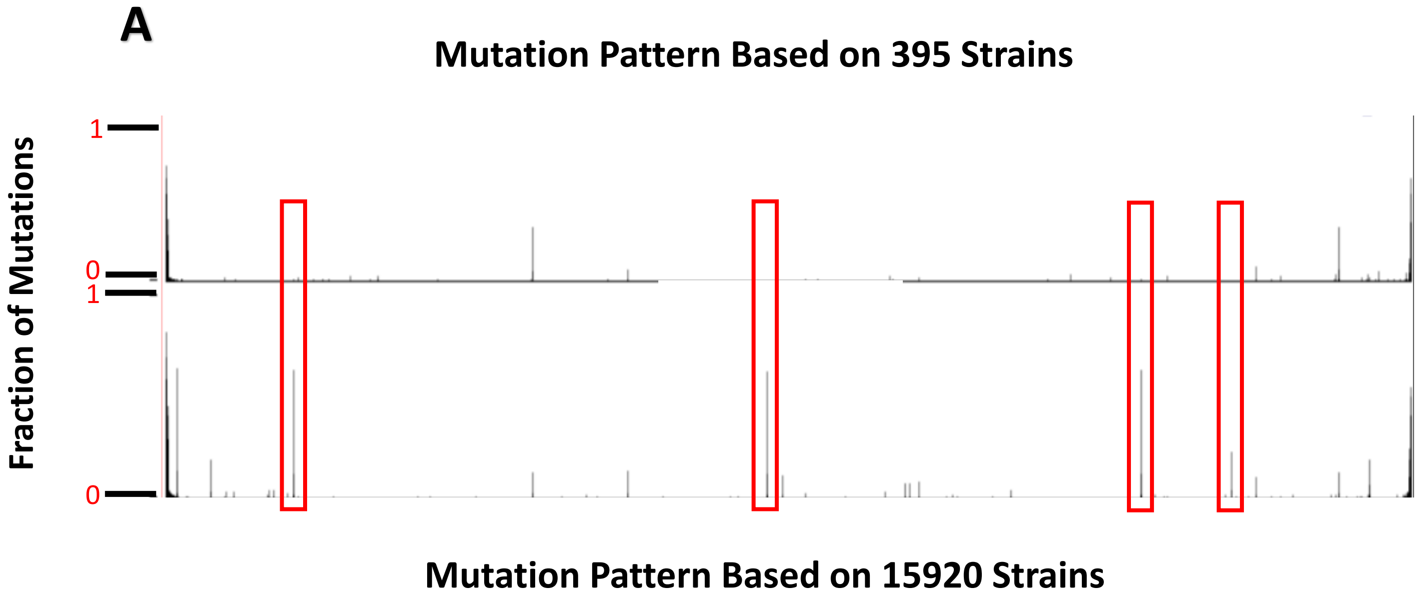


Zhang et al. Figure S1

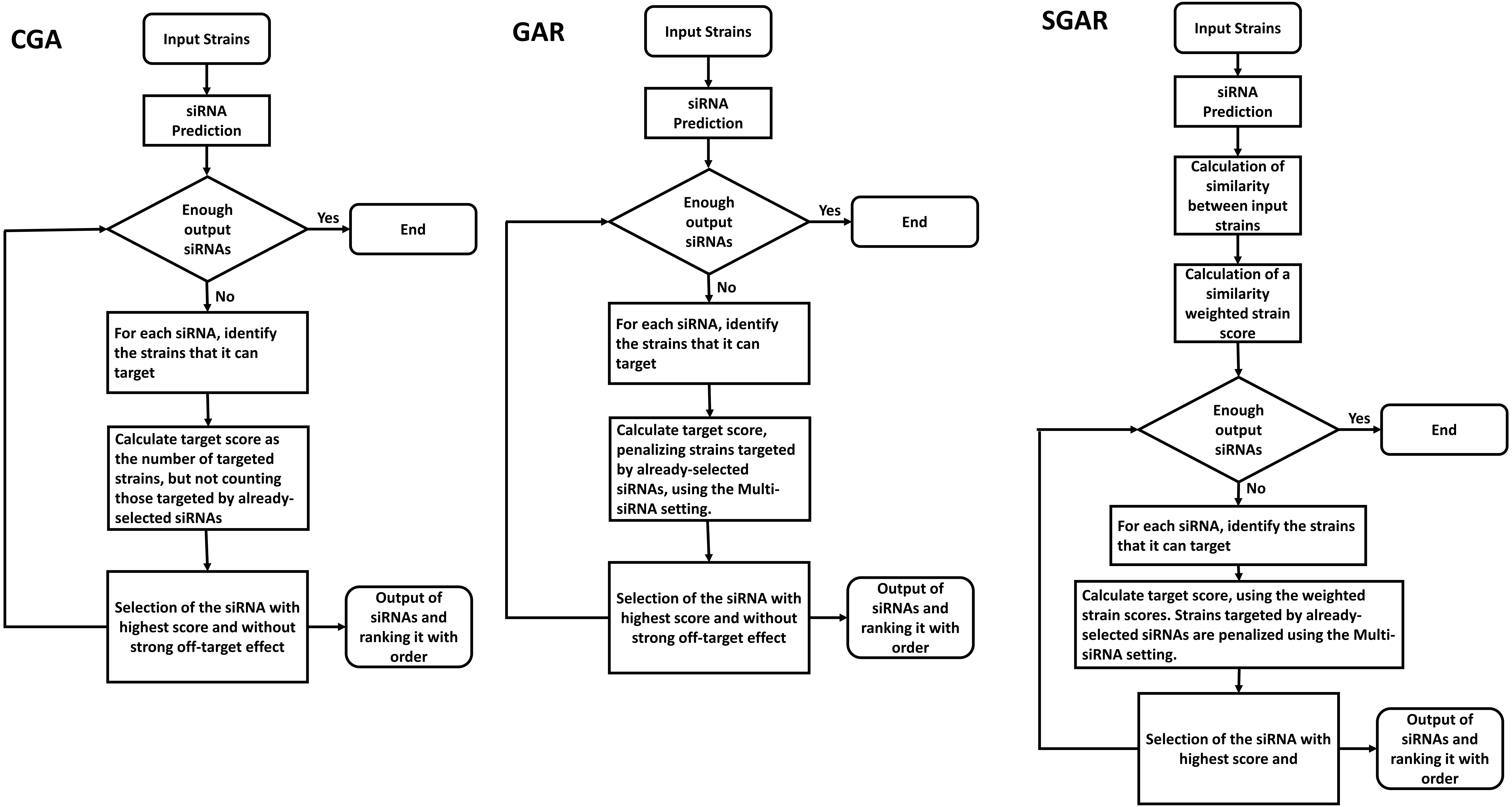


Zhang et al. Figure S2

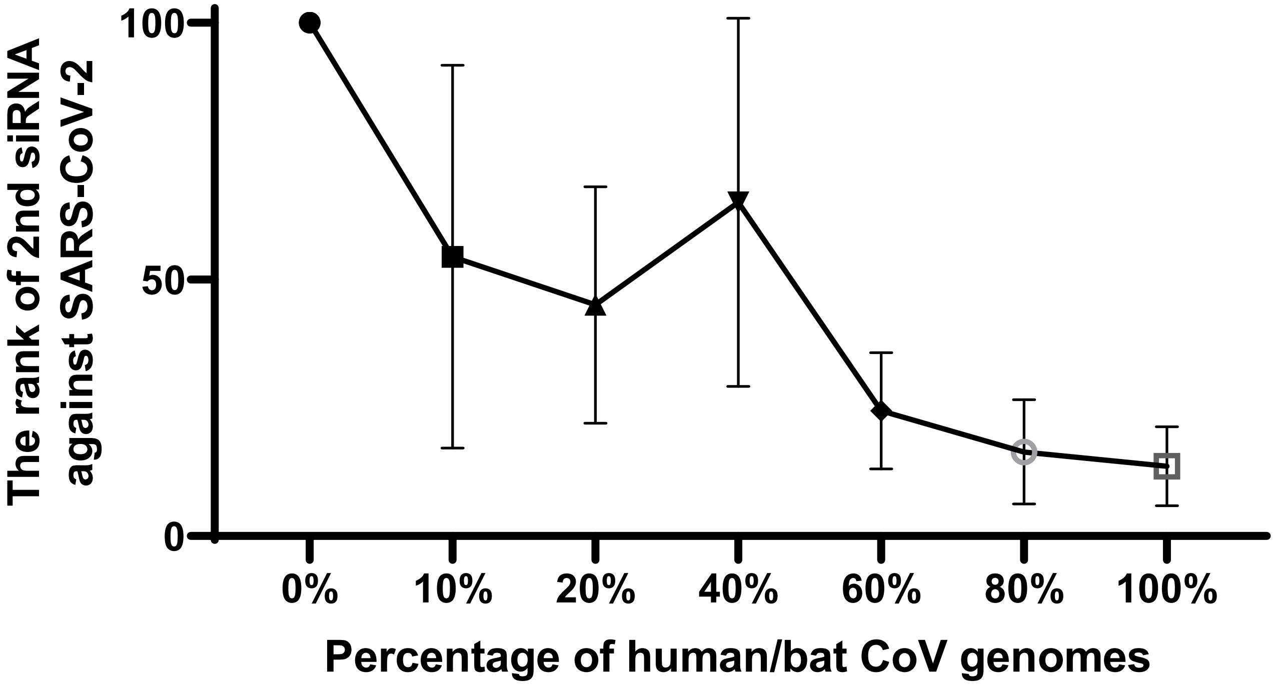


Zhang et al. Figure S3

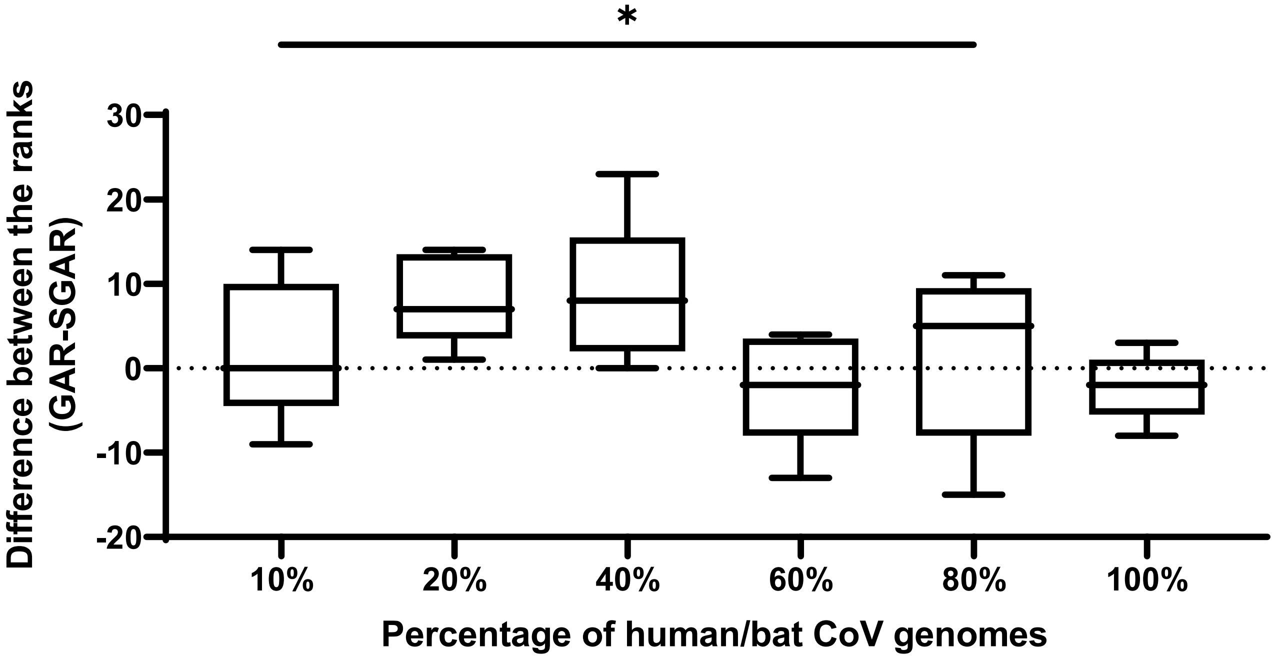


Zhang et al. Figure S4

A

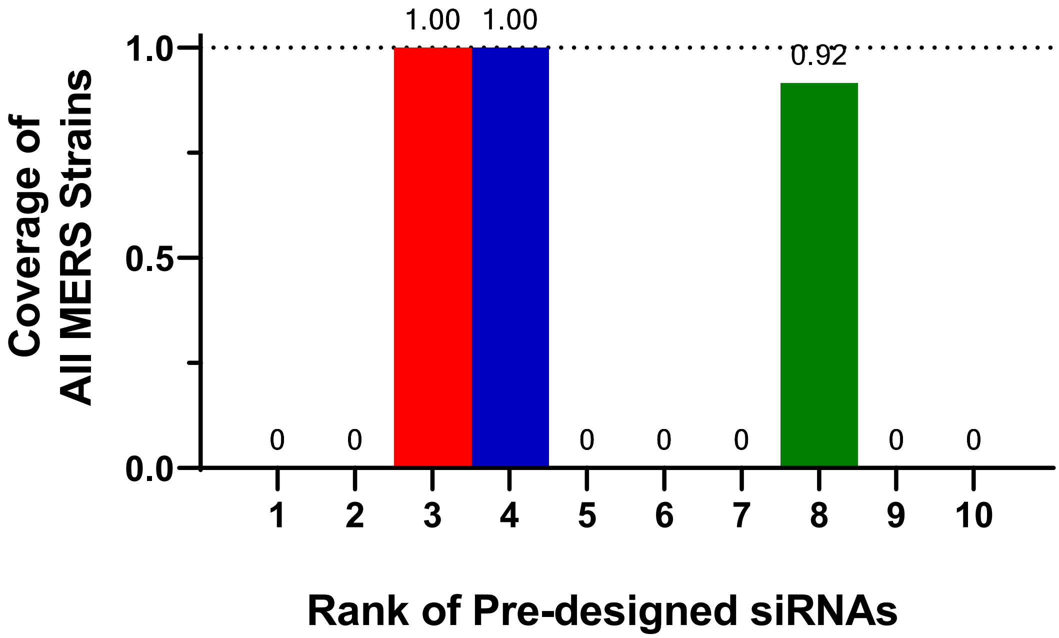


B

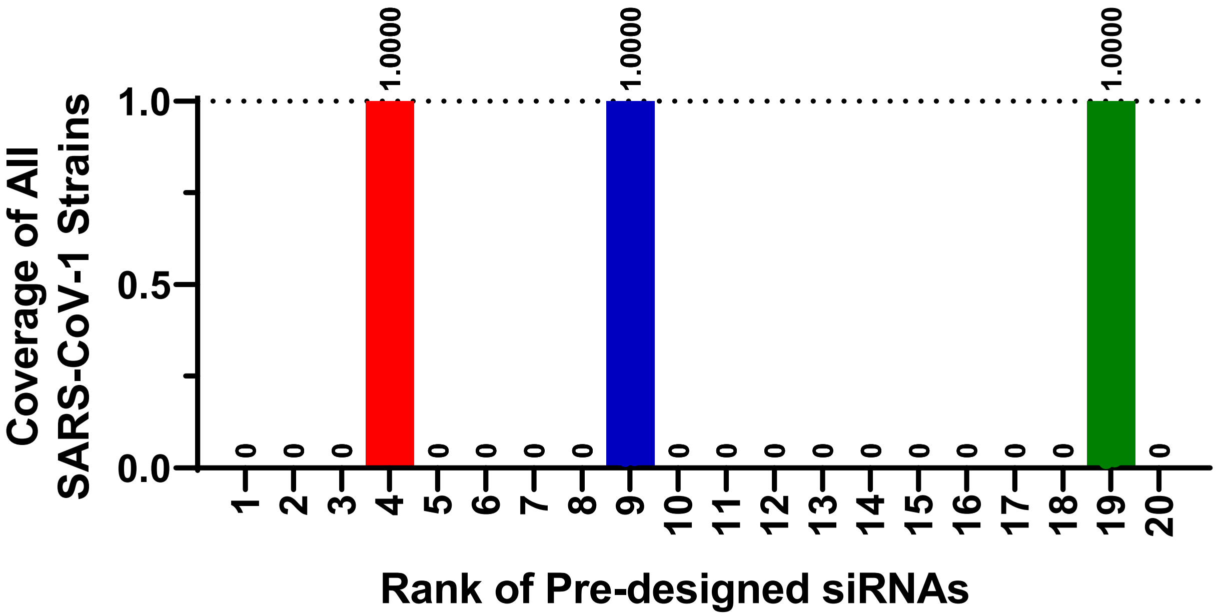


Zhang et al. Figure S5

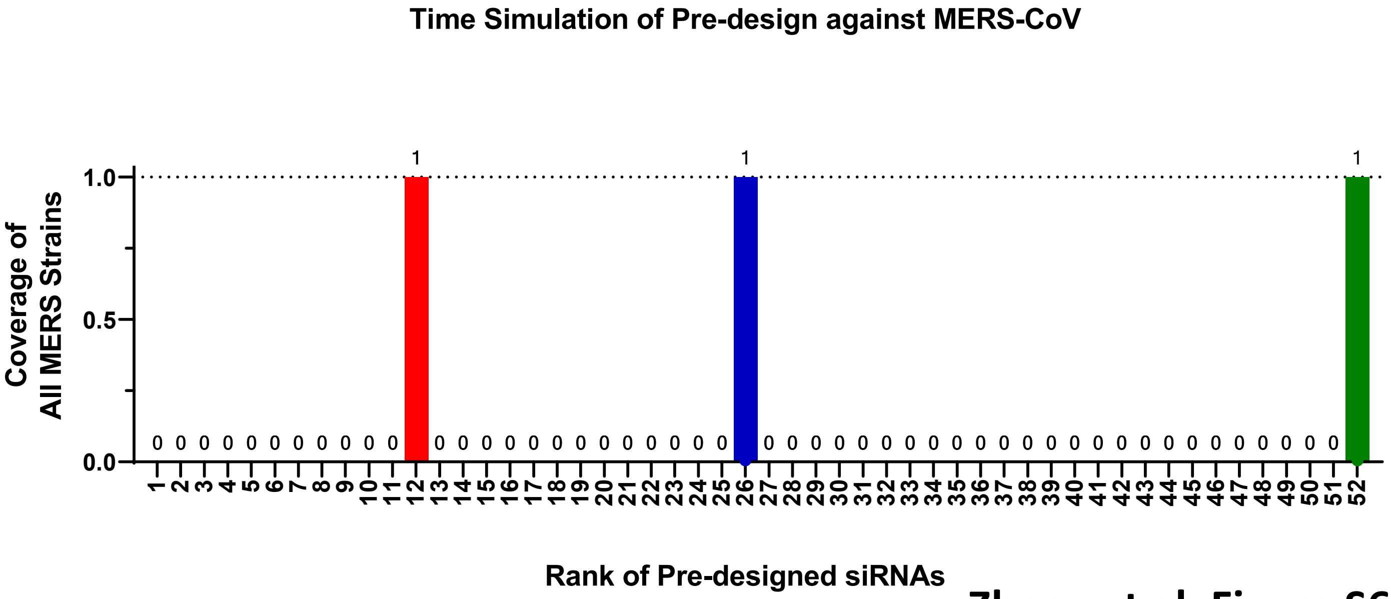


**Zhang et al. Figure S6**
